# SKOR1 mediates FER kinase-dependent invasive growth of breast cancer cells

**DOI:** 10.1101/2022.02.22.481420

**Authors:** Lilian Sluimer, Esme Bullock, Max Rätze, Lotte Enserink, Celine Overbeeke, Marten Hornsveld, Valerie G. Brunton, Patrick W.B. Derksen, Sandra Tavares

**Affiliations:** Department of Pathology, University Medical Center Utrecht, Utrecht, The Netherlands; Edinburgh Cancer Research UK Centre, University of Edinburgh, Edinburgh, Scotland, UK; Department of Molecular Cell Biology, Cancer Genomics Centre Netherlands and Centre for Biomedical Genetics, Leiden University Medical Center, Leiden, The Netherlands; i3S - Instituto de Investigação e Inovação em Saúde, Universidade do Porto, Porto, Portugal; IPATIMUP - Instituto de Patologia e Imunologia Molecular da Universidade do Porto, Porto, Portugal

**Keywords:** Breast cancer, FER tyrosine kinase, SKOR1, metastasis, analogue-sensitive, CRISPR-Cas9

## Abstract

High expression of the tyrosine kinase FER is an independent prognostic factor that correlates with poor survival in breast cancer patients. To investigate whether the kinase activity is essential for FER oncogenic properties, we developed an ATP analogue-sensitive knock-in allele (FER^ASKI^). Specific FER kinase inhibition in MDA-MD-231 cells reduces migration, invasion, and metastasis in a mouse model of breast cancer. Using the FER^ASKI^ system, we identify SKI family transcriptional corepressor 1 (SKOR1) as a direct FER kinase substrate. SKOR1 loss phenocopies FER inhibition, leading to impaired proliferation, migration and invasion, and inhibition of breast cancer growth and metastasis formation in mice. We show that the candidate FER phosphorylation residue, SKOR1-Y234, is essential for FER-dependent tumor progression features. Finally, our work suggests that the SKOR1-Y234 residue promotes Smad2/3 signaling through SKOR1 binding to Smad3 attenuation.

Our study thus identifies SKOR1 as a mediator of FER-dependent breast cancer progression, advocating FER kinase inhibition as a candidate strategy to treat high-grade breast cancers.

**Summary:** The SKI FAMILY TRANSCRIPTIONAL COREPRESSOR 1 (SKOR1) has been mainly associated with neuronal development. Now, Sluimer *et al*. identify SKOR1 as a new substrate of the oncogenic tyrosine kinase FER, and a driver of TNBC progression.

## Introduction

Metastasis underlies the mortality in most high-grade and triple negative breast cancer (TNBC) patients(Kirsch and Loeffler, 2005). Clinical management of these aggressive breast cancer types is challenging due to a lack of responses to standard chemotherapy treatment and hormone receptor antagonist regimens(Aversa et al., 2014; Kennecke et al., 2010). Together, these features underscore an unmet need: the identification of markers that identify drug benefit and/or allow targeted intervention of high-risk breast cancers.

The FPS/FES-related tyrosine kinase FER is a non-receptor tyrosine kinase that was identified as a promising candidate for targeted therapy of metastatic breast cancer(Ivanova et al., 2013). FER kinase regulates cell-cell and cell-matrix contacts, possibly through interaction with adherens junction components p120-catenin and β-catenin(Greer, 2002), by controlling actin polymerization(Ladwein and Rottner, 2008) or through down-regulating the synthesis of glycans that bind to the basement membrane protein laminin(Yoneyama et al., 2012). Moreover, FER controls cell cycle progression and promotes integrin-dependent cell migration and invasion(Ivanova et al., 2013; Kim and Wong, 1995, 1998; Kapus et al., 2000; Arregui et al., 2000). High FER levels have been linked to ovarian, renal and colon cancer progression(Kawakami et al., 2013; Allard et al., 2000; Li et al., 2009; Menges et al., 2010; Takeshima et al., 1998; Zirngibl et al., 2001). In breast cancer, FER expression is an independent predictor of survival, especially in patients negative for lymph-node metastases(Ivanova et al., 2013). Elevated FER levels correlate with high grade breast cancer types such as TNBC and breast cancer brain metastasis(Ivanova et al., 2013; Oshi et al., 2020). Moreover, FER promotes growth and dissemination of melanoma and breast cancer cells in mouse models(Ivanova et al., 2013, 2019). Apart from tyrosine phosphorylation of p120-catenin, β-catenin, actin-binding protein cortactin(Kim and Wong, 1998) and microtubule interactor CRMP2(Zheng et al., 2018), data on direct FER substrates are currently scarce.

Sno/Ski family members have been linked to transforming growth factor β (TGF-β) and bone morphogenic protein (BMP) signaling pathways, possibly through binding and antagonizing Smad proteins(Akiyoshi et al., 1999; Luo, 2004; Luo et al., 1999). Although Sno and Ski are classified as proto-oncogenes, their exact role in tumor progression remains largely unknown. All Sno/Ski family members feature a conserved Sp100/AIRE-1/NucP41/75/DEAF-1 (SAND) motif within their Ski homology domain(Wu et al., 2002). The SAND domain is present in a subset of nuclear proteins that are involved in chromatin-dependent transcriptional regulation(Bottomley et al., 2001). An interaction loop (I-loop) within the SAND domain allows these proteins to bind DNA(Wu et al., 2002; Bottomley et al., 2001). In addition, it has been described that Ski, SnoN and SKOR2 can interact with Smad2/3 through their N-terminal regions and with Smad4 through their SAND domains(Wu et al., 2002).

SKOR1, or functional Smad suppressor element on chromosome 15 (Fussel-15), is the most recently identified member of the Sno/Ski family(Arndt et al., 2007). Human SKOR1 is primarily, but not exclusively, expressed in the central nervous system in the migratory precursors of cerebellar Purkinje cells(Arndt et al., 2007). SKOR1 can act as a transcriptional co-repressor with homeodomain transcription factor Lbx1, thereby regulating the cell fate of dorsal horn interneurons(Mizuhara et al., 2005). Recent evidence implicates a role for SKOR1 in restless leg syndrome and transcriptional regulation of genes involved in neurodevelopment and iron metabolism(Sarayloo et al., 2020). Additionally, activated fibroblasts express SKOR1 during early phases of wound healing, where it promotes fibroblast migration and affects F-actin and focal adhesion distribution(Arndt et al., 2011). In cancer, functions for SKOR1 have remained largely unexplored, except for a recent study observing SKOR1 expression in breast cancer metastasis to the brain(Oshi et al., 2020).

Here, we have used a chemical genetics approach to identify and validate FER substrates and discovered SKOR1 as a direct substrate of the non-receptor tyrosine kinase FER. Our data demonstrate that SKOR1 is essential for tumor growth and invasion, indicating that the tumor promoting function of SKOR1 depends on the candidate FER phosphorylation tyrosine residue Y234.

## Results

### FER kinase activity controls invasion and metastasis formation in TNBC

Previous studies have established FER kinase as a driver of invasion and metastasis formation in high grade and basal-like breast cancers(Ivanova et al., 2013). To study if FER relies on its kinase activity to drive tumor progression, we generated a FER analogue-sensitive (FER-AS) knock-in allele (FER^ASKI^) in MDA-MB-231 (MM231) cells using CRISPR-Cas9 gene editing (SFig. 1A-D). To do so, the gatekeeper residue in the kinase active site of endogenous FER is mutated, which results in an enlarged ATP-binding pocket. Mutation of methionine 637 to alanine in FER confers unique sensitivity to chemically modified derivatives of PP1, the Src-family-selective inhibitor(Bishop et al., 2000, 1998). Knock-in of the gatekeeper mutation (M637A) results in near-endogenous expression levels of FER-AS when compared to MM231 parental cells (SFig. 1E). Importantly, analogue-sensitive kinase inhibition using the PP1 analogue kinase inhibitor 1-(1,1-dimethylethyl)-3-(1-naphthalenylmethyl)-1H-pyrazolo[3,4-d] pyrimidin-4-amine (NM-PP1) does not influence expression of FER-AS (SFig. 1E) but leads to decreased tyrosine phosphorylated on several proteins (SFig. 1F). Whereas no effect is observed in control MM231 cells (SFig. 1G), treatment of FER^ASKI^ cells with 1 μM NM-PP1 leads to a spread and sessile phenotype accompanied by abundant stress fibers and focal adhesion (FA) formation (Fig. 1A and 1B), indicating that this phenotype is a result of specific inhibition of FER kinase activity. Although FER kinase inhibition in FER^ASKI^ interphase cells using NM-PP1 does not significantly affect proliferation (SFig. 1H), it impairs cell motility in 2D (Fig. 1C). Similarly, while proliferation is not affected upon FER kinase inhibition in 3D cultured cells (Fig. 1D and quantified in SFig. 1I), invasion is significantly impaired upon treatment with NM-PP1 (Fig. 1D-E). Combined, these results provide formal evidence that FER regulates tumor cell invasion and migration through its tyrosine kinase activity.

**Figure 1.**
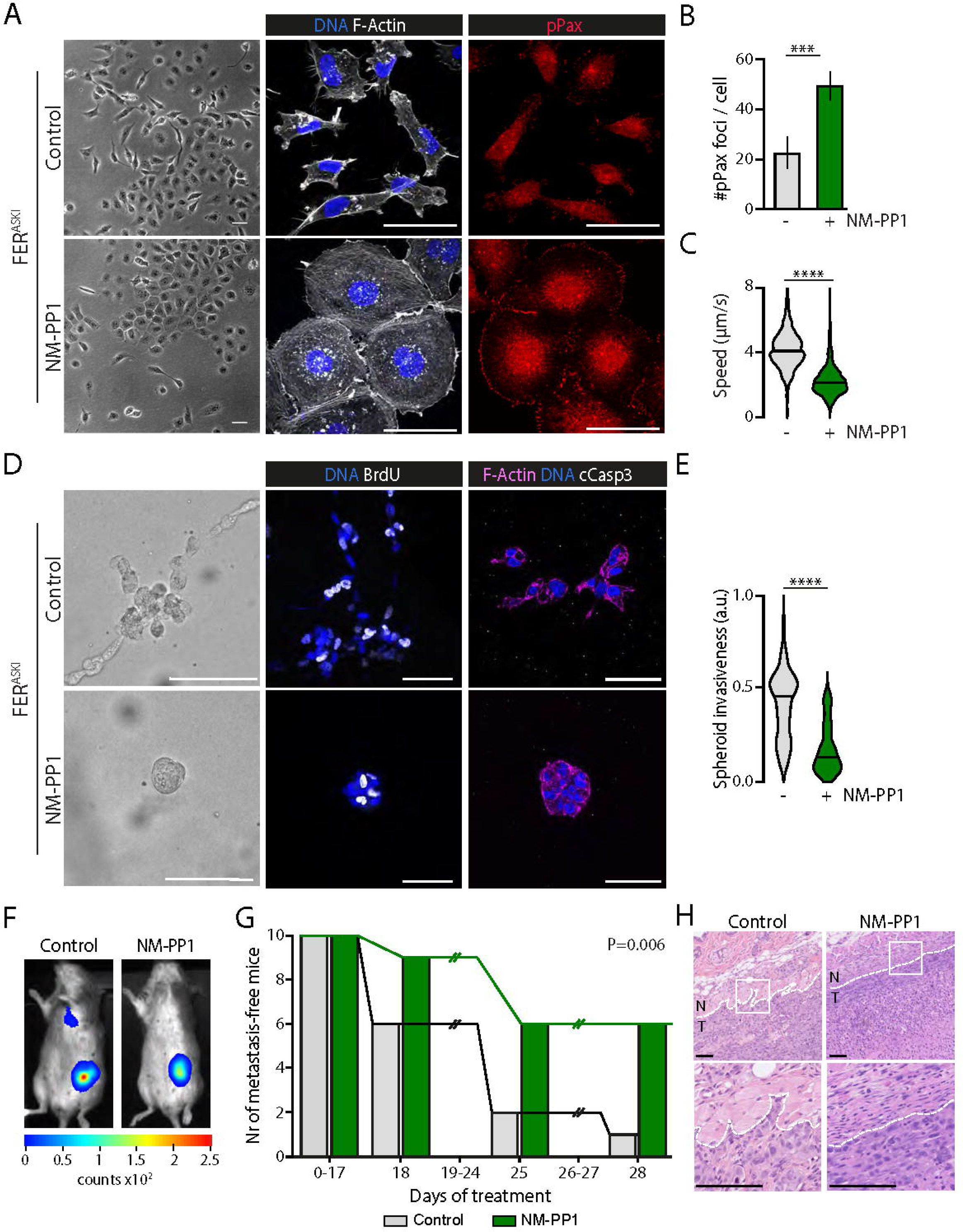
FER kinase activity promotes cell migration and invasion. **A-C**. Inhibition of FER-AS kinase using NM-PP1 induces cell spreading, stress fiber formation and FA formation, and reduces motility in MM231 cells. Cells were imaged by phase-contrast and stained for F-actin (middle panel, white), DNA (DAPI, middle panel, blue) and pPax (right panel, red). Scale bars, 50 μm. Quantifications of pPax foci are shown in (B). Migration speed was quantified in (C) using live fluorescence imaging of MM231 FER^ASKI^ cells treated with control or with NM-PP1. **D-E**. FER kinase is essential for 3D invasion. FER^ASKI^ cells were plated in BME as tumor spheroids, treated with NM-PP1 and imaged by phase-contrast and stained for BrdU incorporation (middle panel, white), cleaved Caspase-3 (right panel, white), F-actin (right panel, magenta) and DNA (DAPI, blue). Scale bars, 50 µm. Quantification of tumor spheroid invasiveness is shown in (E). Error bars denote standard deviation; * P<0.05, ** P<0.01, **** P<0.0001, ns indicates non-significant; one-way ANOVA. **F-G**. Inhibition of FER kinase activity prolongs metastasis-free survival in mice. FER^ASKI^ cells were xenografted, and tumor volume and metastasis formation were monitored upon administration of DMSO (control, N=10) or NM-PP1 (N=10). Metastasis was determined by lung bioluminescence or post-mortem assessment of lung metastatic foci. Representative images of mice treated with control or NM-PP1 at end points are shown in (F). Survival of mice upon treatment is shown in (G). Statistical analysis of survival distributions for the different treatments was performed using Log Rank (Mantel-Cox) test. **H**. FER kinase activity regulates tumor invasion. Dashed white lines indicate tumor (T)-normal (N) breast tissue border. Inset images correspond to a 200% magnification. Scale bars, 100 μm.

Although FER depletion is sufficient to inhibit metastasis formation of MM231 cells in mice(Ivanova et al., 2013), it remained unclear if this was due to inhibition of its tyrosine kinase activity. To test this, we orthotopically transplanted luciferase-tagged FER^ASKI^ cells in recipient RAG2-deficient mice and started treatment with NM-PP1 upon formation of palpable primary tumors (∼ 50 mm^3^). Tumor growth was monitored longitudinally, and mice were sacrificed once lung metastases were detected by bioluminescence imaging. We do not observe differences in primary tumor volumes between the treated and untreated cohorts (Fig 1F and quantified in SFig. 1J), in agreement with published findings showing that loss of FER function *in vivo* mainly affects invasion and metastasis(Ivanova et al., 2013). Untreated animals show a median metastasis-free latency of approximately 22 days (Fig. 1G), supporting published findings using FER loss(Ivanova et al., 2013) and showing that the FER^ASKI^ allele is fully functional *in vivo*. In contrast, inhibition of FER kinase activity with NM-PP1 delays the median development of metastasis to 28 days when compared to the control group (p=0.006; Fig. 1G). NM-PP1-treated mice develop tumors that show reduced local invasion in the surrounding tissue compared to control mice (Fig. 1H). Together, these results demonstrate that FER kinase activity promotes tumor invasion and metastasis of TNBC cells in mice.

To analyze the impact of FER on phosphorylated downstream effectors, we used a Doxycycline (Dox)-inducible FER knock-down (FERiKD) combined with a reverse-phase protein array (RPPA) analysis (SFig. 2A and SFig.3A). Expression and phosphorylation were assayed using stable FERiKD in two high grade basal breast cancer cell types, MM231 (SFig. 2B and Table S1) and SUM149PT (SFig. 3B and Table S2). Dox-induced loss of FER expression attenuates the activation of multiple key signaling pathways in MM231 cells (Table S1), such as Rb phosphorylation at Ser807 and Ser780 (SFig. 2C), sites required for G0-G1 transition(Ren and Rollins, 2004). As loss of FER was shown to inhibit cell cycle progression and impair proliferation of MM231 cells(Ivanova et al., 2013; Pasder et al., 2006), our data further confirm a crucial role for FER in regulating TNBC cell proliferation. In line with the kinase-specific inhibition of FER (SFig. 1G), we found that EGFR Tyr1173 phosphorylation is reduced upon loss of FER in MM231 cells (Table S1), confirming EGFR as a FER downstream target(Guo and Stark, 2011). Moreover, we observe altered levels of VEGFR-Tyr1175, PDGFR-Tyr1021 and IGF1R-Tyr1162/3 upon FER loss (Table S1 and Table S2), suggesting that FER indirectly controls the function and/or processing of multiple growth factor receptors. Interestingly, Smad2/3 phosphorylation levels are consistently decreased after FER loss on several Serine epitopes in both cell systems used (Table S1 and Table S2), which suggests a role for the regulation of TGFβ/BMP signals by FER.

### SKOR1 is a direct FER substrate that promotes migration and invasion

To identify novel pathways regulating TGFβ/BMP signals downstream of FER, we focused our efforts exploring a less studied member of the Ski/SnoN family of TGFβ/BMP signaling repressors, SKOR1. We selected SKOR1 because it has been linked to the regulation of BMP signaling through interaction with Smads(Arndt et al., 2007; Fischer et al., 2012), its expression has been observed in breast cancer brain metastases(Oshi et al., 2020), and its role in tumor progression remains largely unknown. We validated SKOR1 as a direct FER substrate *in vitro* using a GST-tagged SKOR1 and ATP incorporation (Fig. 2A-B). SKOR1 phosphorylation is specific, because either using a kinase-dead FER (D742R) or pharmacological inhibition of the gatekeeper FER kinase using NM-PP1 significantly inhibits SKOR1 phosphorylation (Fig. 2A-B). Moreover, phosphorylation is reduced when using SKOR1-Y^NULL^, a SKOR1 mutant in which all tyrosine residues in exon 5 are mutated to alanine (SFig. 4A), indicating that FER-dependent phosphorylation occurs specifically on tyrosine residues.

**Figure 2.**
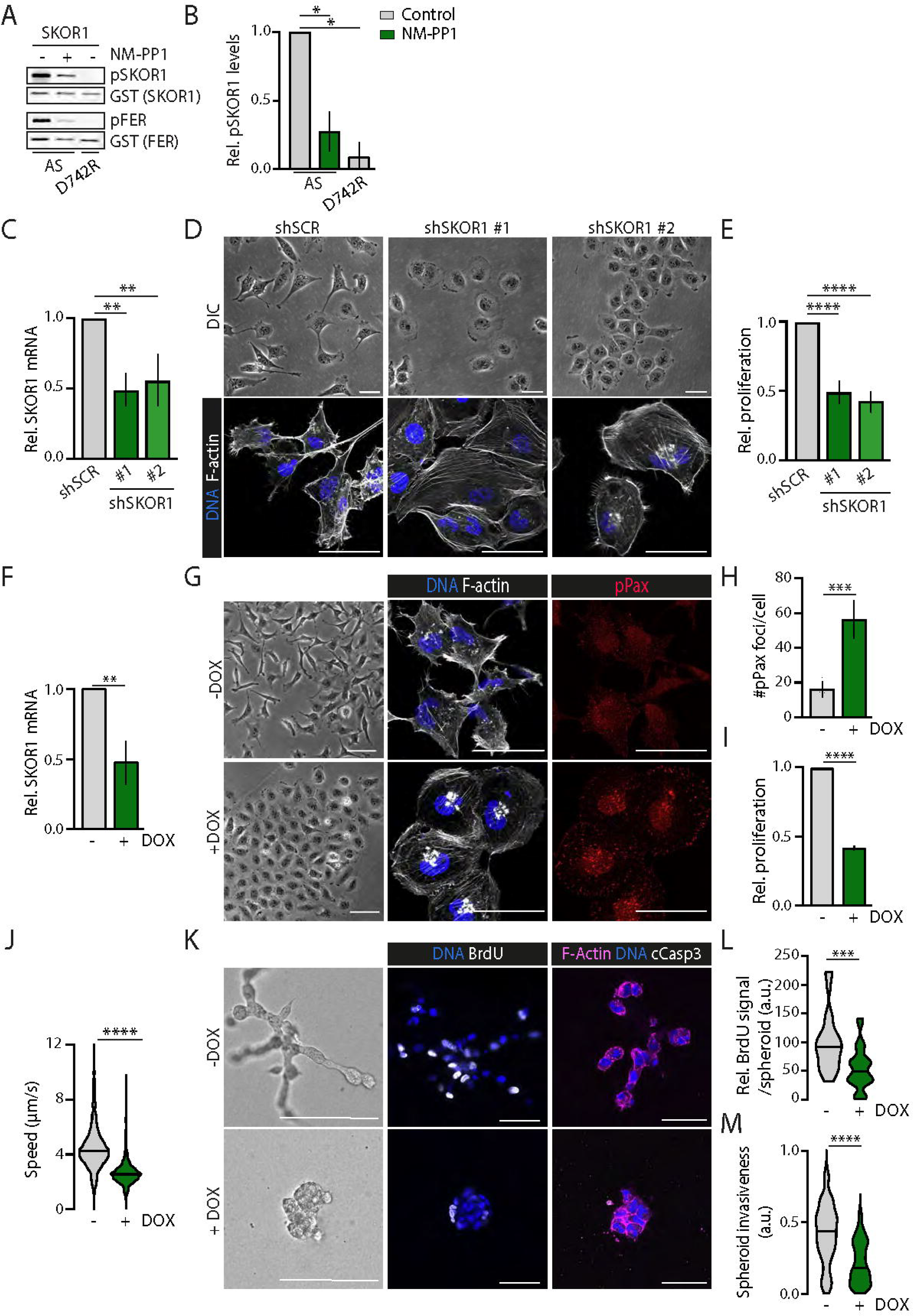
SKOR1 is a FER kinase substrate that promotes breast cancer cell proliferation and invasion. **A-B**. FER phosphorylates SKOR1 in vitro. Kinase assay using recombinant GST-analog sensitive FER (AS) or GST-kinase dead FER (D742R) in the presence of N-benzyl(Bn)-ATPγS, with (+) or without (-) NM-PP1. GST-tagged SKOR1^WT^ was used as substrate. Auto-thio-phosphorylation of FER serves as positive control. GST-FER was used as loading control. Quantifications are from at least three independent experiments (B). **C-D and F-H**. SKOR1 is critical for TNBC cell morphology in 2D. Validation of SKOR1 knock-down using real-time quantitative-PCR amplification in stable (C) and Dox-inducible (F) KD cells. MM231 stable SKOR1 KD cells were imaged using phase-contrast (upper panel) and stained for F-actin (lower panel, white) and DNA (DAPI, blue) (D). MM231 SKOR1iKD cells were treated with Dox and imaged by phase-contrast (left panel) and stained for F-actin (middle panel, white), pPax (right panel, red) and DNA (DAPI, blue) (G). Scale bars, 50 µm. Quantifications of pPax foci are shown in (H). Note the cell spreading, stress fiber formation and increase in FA formation upon SKOR1 loss. **E and I**. SKOR1 controls TNBC cell proliferation. Shown are quantifications of colony formation assays of stable (E) and Dox-inducible (I) SKOR1 KD cells. **J**. SKOR1 is important for 2D cell motility. Migration speed was quantified in using live fluorescence imaging of MM231 SKOR1iKD cells transfected with pGK-GFP -/+ Dox grown in 2D. **K-M** SKOR1 is required for proliferation and invasion in 3D. MM231 SKOR1iKD cells were treated with Dox and plated in BME to form 3D spheres. Cells were imaged by phase-contrast and stained for BrdU (middle panel, white), cleaved Caspase-3 (right panel, white), F-actin (right panel, magenta) and DNA (DAPI, blue). Scale bars, 50 µm. Quantifications of BrdU positive nuclei and spheroid invasion are shown in (L) and (M), respectively. Error bars denote SD; * P<0.05, ** P<0.01, **** P<0.0001, ns indicates non-significant; one-way ANOVA.

To analyze the role of SKOR1 in high grade metastatic breast cancer cells, we performed a stable knock-down of SKOR1 in MM231 using two independent shRNA hairpins (Fig. 2C-E). SKOR1 mRNA levels were stably and consistently reduced to approximately 50% in multiple independent experiments (Fig. 2C). Reducing SKOR1 levels using either hairpin results in a stark reduction in cell numbers compared to the scrambled control shRNA (compare day 1 to day 4, SFig. 4B), suggesting that SKOR1 is necessary for cell proliferation and survival. After stable integration and selection, we could confirm that SKOR1-depleted cells display a decreased proliferative capacity (Fig. 2E). Strikingly, loss of SKOR1 phenocopies FER depletion(Ivanova et al., 2013) and inhibition (Fig. 1A and Fig. 2D), including extensive cell spreading and collective growth as non-motile cells. SKOR1 depletion also induces loss of lamellipodia and F-actin stress fiber formation in 2D (Fig. 2D), a characteristic feature of FER loss or inhibition in MM231 cells ((Ivanova et al., 2013) and Fig. 1A).

To temporally and reversibly control the SKOR1 knock-down, we employed a Dox-inducible lentiviral shRNA system, which reproducibly results in a 50% reduction of SKOR1 expression after induction (SKOR-iKD; Fig. 2F). Inducible SKOR1 loss results in extensive cell spreading, formation of stress fibers (Fig. 2G) and a significant increase in FA formation (Fig. 2H). In addition, we observed that SKOR1 knock-down leads to impaired 2D proliferation (Fig 2I) and impairment of single cell migration when compared to cells cultured in control conditions (Fig. 2J).

We next cultured SKOR1iKD cells in 3D basement membrane extract (BME) gels and observed that SKOR1 depletion fully prevents invasion when compared to control spheroids that exhibit highly branched, invasive and disorganized colonies (Fig. 2K and 2M). Because SKOR1-depleted cells show clear growth defects when cultured in 2D, we assessed proliferation and apoptosis using BrdU incorporation and cleaved caspase-3 as markers, respectively. These analyses indicate that the reduction in colony formation upon SKOR1 depletion is mainly due to an impairment in proliferation in 3D (Fig. 2L), because we observed no differences in apoptosis compared to controls (Fig. 2K).

### SKOR1 tyrosine 234 is required for cellular proliferation and invasion

Our chemical genetics and proteomics studies had identified SKOR1-Tyrosine 234 (Y234) as a candidate FER phosphorylation site, a residue that resides in the I-loop of the SAND domain (Fig. 3A)(Arndt et al., 2007). We therefore stably expressed a GFP-tagged and RNAi-resistant wild-type (WT) SKOR1 cDNA, or a SKOR cDNA harboring a phenylalanine at position 234 (Y234F) in MM231 SKOR1iKD cells (SKOR1^RECON^; Fig. 3B), which prevents phosphorylation of SKOR1-Y234 without altering the protein structure. Upon SKOR1 reconstitution, we observed that the WT cDNA fully rescues the SKOR1 knock-down phenotype, presenting a migratory and spindle-like morphology resembling the parental MM231 morphology (Fig. 3C, upper panels). Conversely, SKOR1^RECON^::Y234F cells fail to migrate and exhibit a spread and sessile phenotype (Fig. 3C, bottom panels). These results coincide with the formation of stress fibers and a marked increase in the number of FA sites (Fig. 3C, quantified in Fig. 3D), which suggests that the SKOR1 Y234 residue is involved in the regulation of F-actin dynamics and FA formation in MM231 cells. SKOR1^WT^ and SKOR1^Y234F^ localize throughout the cytosol in vesicular-like structures that are most prominent in the perinuclear region (Fig. 3C). Interestingly, reconstitution with SKOR1^Y234F^ results in a 60% decrease in cellular proliferation in 2D compared to SKOR1^WT^ (Fig. 3E). Next, we assessed 3D cancer cell invasion, proliferation and apoptosis. We unexpectedly observe that the SKOR1-Y234 residue does not significantly contribute to cellular proliferation, nor apoptosis in this setting (Fig. 3F and 3G). However, the SKOR1^Y234F^ mutant is unable to rescue invasion of MM231 cells in 3D (Fig. 3F, quantified in Fig. 3H). Together, our results show that SKOR1-Y234, a FER kinase candidate substrate residue, is necessary for the invasion of the triple negative MM231 cells.

**Figure 3.**
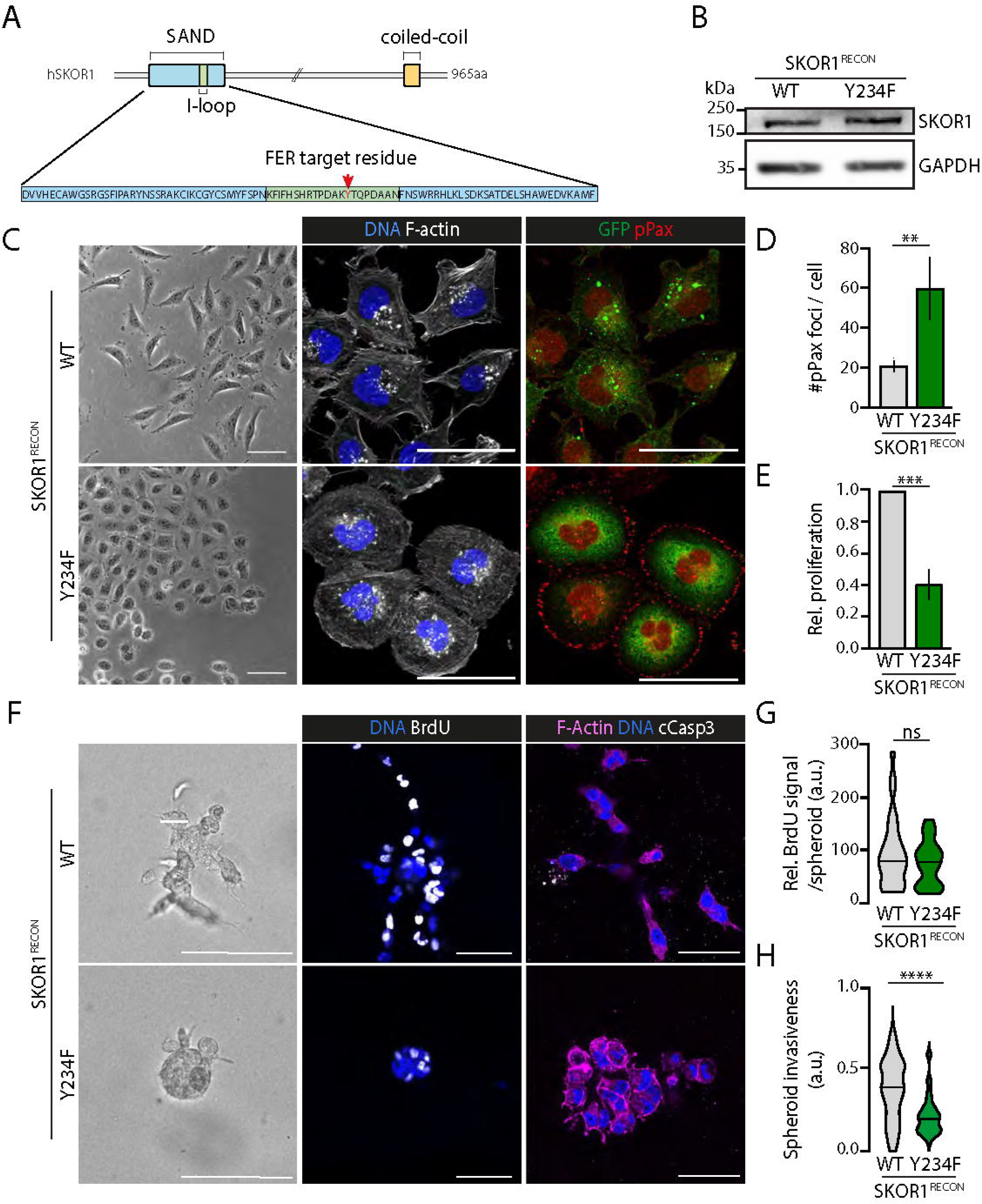
SKOR1-Y234 is required for cell motility and invasion. **A**. The FER target residue of SKOR1 resides in the I-loop of the SAND domain. Schematic of human SKOR1 domains. **B-H**. MM231 SKOR1iKD cells were reconstituted (SKOR1^RECON^) with either SKOR1 wild-type (SKOR1^WT^) or SKOR1 mutant whereby tyrosine 234 was substituted for phenylalanine (SKOR1^Y234F^). **B**. SKOR1^WT^ and SKOR1^Y234F^ are equally expressed. Shown are protein levels of SKOR1 in indicated cells as determined by western blotting. GAPDH was used as loading control. **C-E**. The SKOR1 Y234 residue is required for motility, invasion, and proliferation. Cells were stained for F-actin (middle panel, white), pPax (right panel, red) and DNA (DAPI, blue) (C). Scale bars, 50 µm. Note the extensive cell spreading and increase in the number of pPax foci in SKOR1^Y234F^ cells. Note the cytoplasmic localization of SKOR1 in the WT and Y234F reconstituted cells (right panel). Quantifications of FA formation shown in (D). Colony formation assays were performed and quantified (E). **F-H**. The Y234 residue regulates 3D invasion. SKOR1^WT^ or SKOR1^Y234F^ reconstituted cells were plated in BME as tumor spheroids. Cells were imaged by phase-contrast and stained for BrdU incorporation (middle panel, white), cleaved Caspase-3 (right panel, white), F-actin (right panel, magenta) and DNA (DAPI, blue). Scale bars, 50 µm. Quantifications of BrdU positive nuclei and tumor spheroid invasiveness are shown in (G) and (H), respectively. Error bars denote SD; * P<0.05, ** P<0.01, **** P<0.0001, ns indicates non-significant; one-way ANOVA.

Because of the overlap in phenotype when inhibiting FER or depleting SKOR1, we used the FER^ASKI^ model and transduced SKOR1^WT^ or SKOR1^Y234F^ to assess functional inter-dependence. Control FER^ASKI^::SKOR1^WT^ cells display a highly invasive phenotype, with lamellipodia formation and spindle-like shaped cells (Fig. 4A). SKOR1^WT^ fully prevents the phenotypical consequences of FER kinase inactivation in the presence of NM-PP1, sustaining a spindle-like and motile cell phenotype with sparse FA sites (Fig. 4A, bottom panels and Fig. 4B). Control FER^ASKI^::SKOR1^Y234F^ cells exhibit a mixed/hypomorphic phenotype, whereby migratory cells coincide with cells that show extensive cell spreading and an increase in FA formation (Fig. 4A and 4B). Inhibition of FER kinase activity using NM-PP1 induces a further increase in cell spreading and an accompanying transition from a spindle-like cell shape to a spread phenotype with an increased number of FA sites in FER^ASKI^::SKOR1^Y234F^ cells (Fig. 4A and 4B). Whereas inhibition of FER kinase alone does not affect cell proliferation in 2D (SFig. 1I), proliferation defects are observed in FER^ASKI^::SKOR1^Y234F^ cells treated with NM-PP1 (Fig. 4C), indicating that SKOR1-Y234 is involved in TNBC cell proliferation.

**Figure 4.**
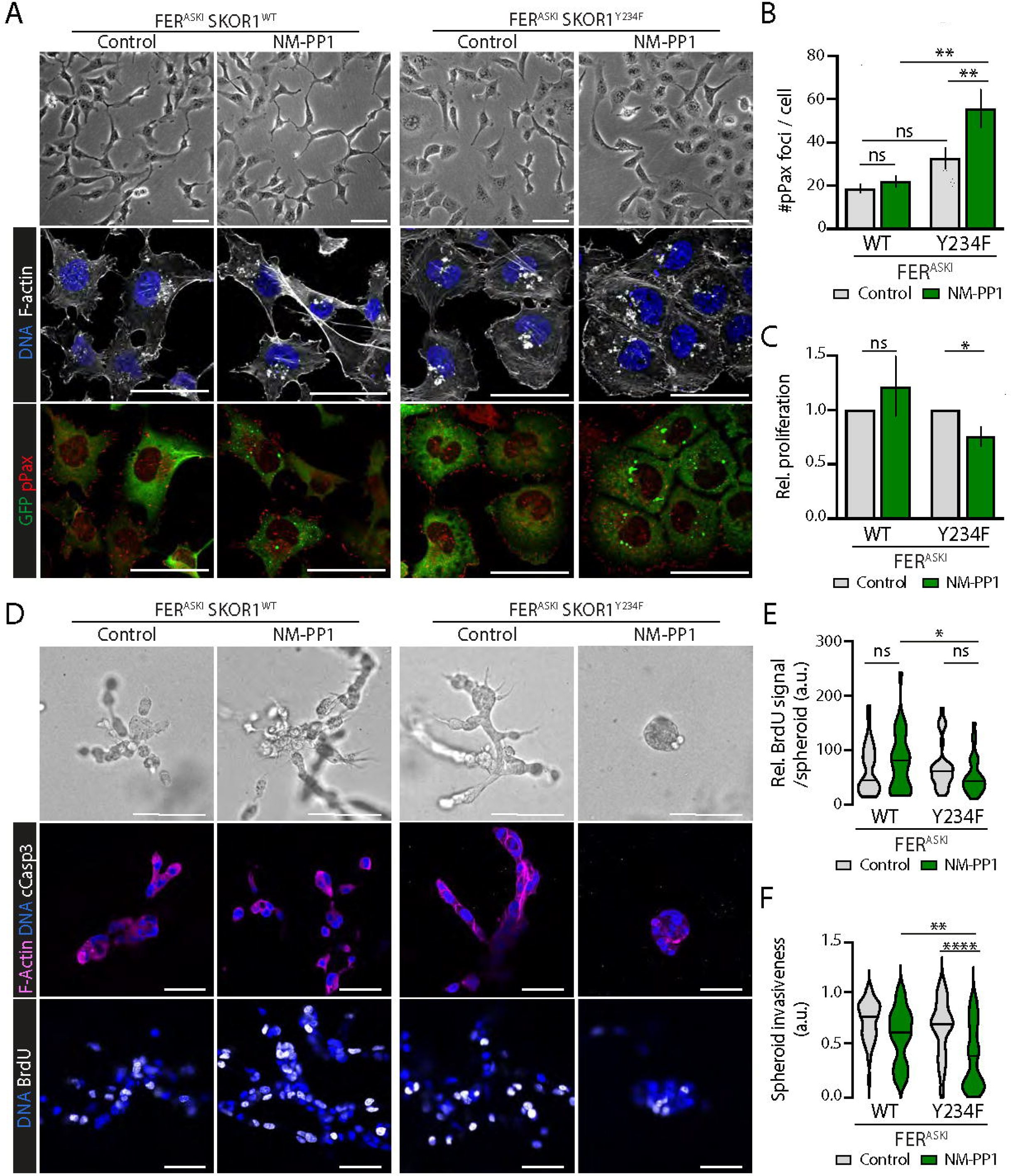
SKOR1 promotes FER-induced migration and invasion through its Tyrosine 234 residue. **A-C**. SKOR1-Y234 controls FER-induced cell motility and proliferation. MM231 FER^ASKI^ cells expressing SKOR1^WT^ or SKOR1^Y234F^ (green) were imaged by phase-contrast and stained for F-actin (middle panel, white), pPax (lower panel, red) and DNA (DAPI). Scale bars, 50 µm. Quantifications of pPax foci are shown in (B). Colony formation assays were performed and quantified (C). **D-F**. SKOR1-Y234 is critical for FER-induced 3D invasion. MM231 FER^ASKI^ cells expressing SKOR1^WT^ or SKOR1^Y234F^ (green) were cultured in the presence of NM-PP1 and plated in BME as tumor spheroids. Cells were imaged by phase-contrast and stained for BrdU incorporation (middle panel, white), cleaved Caspase-3 (right panel, white), F-actin (right panel, magenta) and DNA (DAPI, blue). Scale bars, 50 µm. Quantifications of BrdU positive nuclei and tumor spheroid invasiveness are shown in (E) and (F), respectively. Error bars denote SD; * P<0.05, ** P<0.01, **** P<0.0001, ns indicates non-significant; one-way ANOVA.

We also studied the effect of the SKOR1-Y234F residue on 3D invasion in the context of FER kinase function and observed that SKOR1^Y234F^, in contrast to SKOR1^WT^, is unable to rescue the impaired invasive growth caused by FER inactivation (Fig. 4D). Treatment with NM-PP1 leads to a full inhibition of both invasion and proliferation in FER^ASKI^::SKOR1^Y234F^ cells, whereas FER^ASKI^::SKOR1^WT^ cells form highly invasive structures upon NM-PP1 treatment (Fig. 4D-F). These data suggest that the SKOR1^Y234^ residue functions downstream of FER kinase to promote tumor cell proliferation and invasion of high-grade breast cancer cells.

### SKOR1 regulates BMP/TGFβ signaling and binds Smad2/3 in TNBC cells

We next tested if SKOR1-Y234 modulation leads to phosphorylation changes in signaling pathways. For this we used a reverse phase protein array (RPPA) combined with SKOR1 reconstitution MM231 SKOR1iKD cells (Fig. 5A and Table S3). Reconstitution with SKOR1^Y234F^ reduces phosphorylation of EGFR signaling and its downstream effectors such as 4EBP1 and MAP kinase on multiple epitopes (Fig. 5B, right panel, and Table S3). When comparing SKOR1^Y234F^ cells to FERiKD cells, we observed that many substrates altered by SKOR1^Y234F^ expression are similarly affected by FER depletion, including Rb (Ser780, Ser807), mTOR and EGFR (Tyr1173) (Fig. 5B). Importantly, like in FER-depleted cells, SKOR1^Y234F^ reconstitution leads to a decrease in Smad2/3 phosphorylation (Ser423, Ser425, Ser467/Ser425) (Fig. 5B), which is in concordance with reported links of SKOR1 to BMP/TGFβ signaling(Arndt et al., 2007; Fischer et al., 2012; Takaesu et al., 2012). In sum, we conclude that both FER depletion and SKOR1^Y234F^ reconstitution result in reduced Smad signaling, supporting our hypothesis that FER regulates Smad signals via SKOR1-Y234.

**Figure 5.**
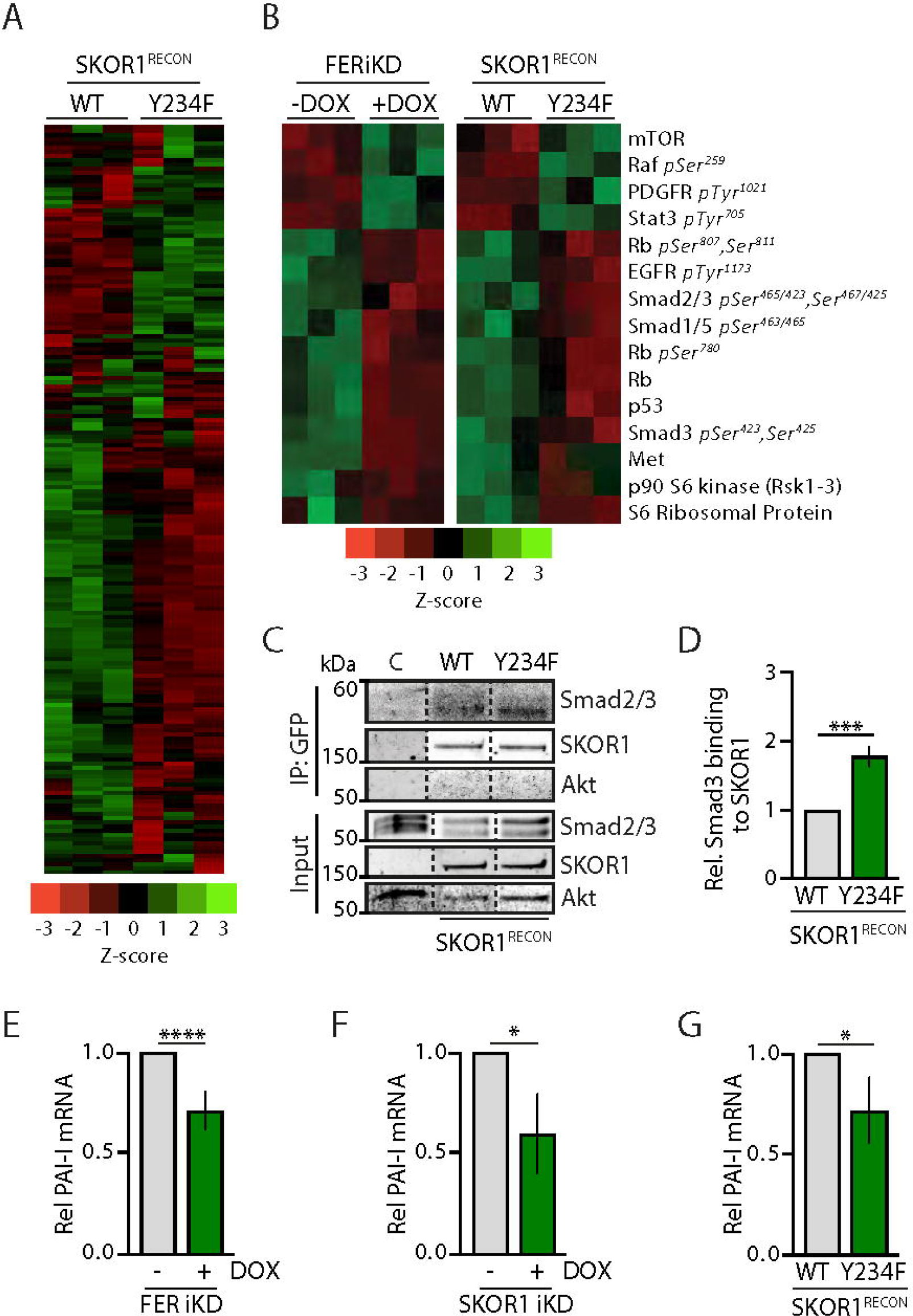
FER and SKOR1-Y234 promote Smad2/3 phosphorylation. **A**. SKOR1-Y234 controls multiple key signaling pathways in MM231 cells. **B**. FER and SKOR1-Y234 similarly induce phosphorylation of Smad2/3 and other key growth signaling effectors. (A and B) Heatmaps representing altered levels of phosphorylated protein residues (Z-scores). Rows represent phosphorylation sites and colors represent magnitude of intensity. The red-to-green scale indicates signal intensity. **C-D**. The SKOR1 Y234 residue controls SKOR1 binding to Smad3. Smad2/3 binding to SKOR1 was assessed in MM231 SKOR1^RECON^ cells using coIP for GFP and western blotting. A parental MM231 cell lysate was used as non-specific IP control. Shown are protein levels of Smad2/3 and SKOR1. Akt was used as loading control. Quantification of Smad2/3 levels bound to GFP-SKOR1^WT^ and GFP-SKOR1^Y234F^ are shown in (D). **E-G**. SKOR1 and FER regulate mRNA expression of PAI-I, a TGFβ target gene. Relative PAI-I expression was determined upon FER depletion (E), SKOR1 depletion (F) and SKOR1^Y234F^ reconstitution (G) in MM231 cells. Quantifications are from at least three independent experiments. Expression was normalized by GAPDH. Error bars denote SD; * P<0.05, **** P<0.0001; t-test.

To verify possible ties between SKOR1 and Smad signaling, we assessed SKOR1 binding to Smad. For this, we performed co-immunoprecipitation (coIP) assays in SKOR1^RECON^ cells, which confirmed that SKOR1^WT^ precipitates with Smad2 and Smad3, whereby we noted a higher affinity to Smad3 (Fig. 5C). SKOR1^Y234F^ reconstitution increases binding to Smad3 compared to SKOR1^WT^ (Fig. 5C, quantified in Fig. 5D), suggesting that phosphorylation of Y234 in SKOR1 weakens binding to Smad3. To further assess if FER and SKOR1 impact Smad-dependent signaling pathways, we assessed the mRNA levels of plasminogen activator inhibitor 1 (PAI-I), a Smad3 effector that is specifically upregulated in Smad3-driven tumor progression(Petersen et al., 2010). Either FER or SKOR1 loss leads to a decrease in PAI-I mRNA expression in MM231 cells (Fig. 5E and F, respectively). We also observe that PAI-I expression is significantly decreased upon SKOR1^Y234F^ reconstitution (Fig. 5G). Together, these data suggest that FER-dependent phosphorylation of SKOR1 on Y234 may regulate binding of SKOR1 to Smad3, thereby facilitating phosphorylation and activation of Smad-dependent signals.

### SKOR1 promotes tumor growth and metastasis in mouse xenografts

We next orthotopically transplanted luciferase-expressing MM231 SKOR1iKD cells in recipient female mice and measured primary tumor growth and metastasis development over time. As SKOR1 loss inhibits proliferation in MM231 cells, we transplanted untreated cells, monitored animals until palpable tumors (∼50 mm^3^) formed in both groups and started Dox administration to induce SKOR1 knock-down. Real-time qPCR was used to quantify SKOR1 mRNA in tumor samples, which confirms our *in vitro* data that SKOR1 expression is downregulated to approximately 50% after knock-down when compared with controls (Fig. 6A). In contrast to FER kinase inhibition, SKOR1 loss significantly affects primary tumor growth (Fig. 6B). Moreover, bioluminescence imaging showed that SKOR1 loss impairs the development of lung metastases, leading to an increased metastasis-free survival (Fig. 6C and 6D). In contrast to control tumors, SKOR1-depleted tumors tend to show expansive growth patterns with little to no invasion into the stroma and adjacent muscle layers of the mammary fat pad (Fig. 6E). From these results we conclude that SKOR1 promotes tumor growth and metastasis of high grade and basal-like breast cancer cells in mice, suggesting that SKOR1 contributes to tumor progression in breast cancer.

**Figure 6.**
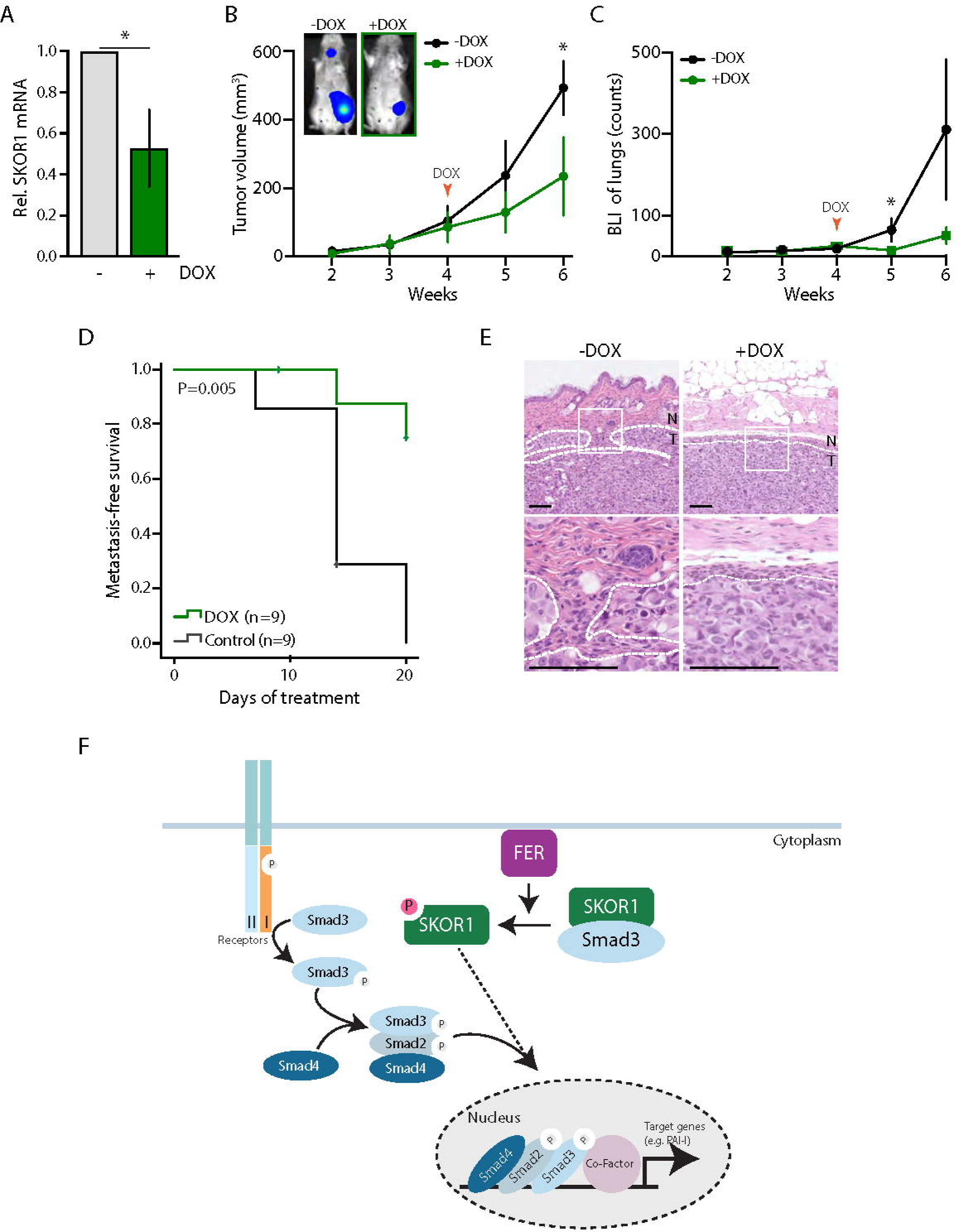
SKOR1 promotes breast tumor growth and metastasis formation. **A**. Validation of SKOR1 knock-down in mice breast tumors by real-time quantitative PCR. **B-D**. SKOR1 promotes breast tumor growth and metastasis formation in mice. Luciferase-expressing MM231 SKOR1iKD cells were xenografted and upon formation of palpable tumors, mice were switched to a Dox-containing diet (red arrow). Primary tumor volume (B) and lung metastasis formation (C) were monitored weekly. Kaplan-Meijer analysis was used to determine cumulative survival of mice treated with (n=9) or without Dox (n=9) (D). **E**. SKOR1 promotes tumor invasion in mice. Representative H&E images of mice treated or untreated with Dox at end points. Dashed white lines indicate tumor (T)-normal (N) breast tissue border. Inset images correspond to a 200% magnification. Scale bars, 100 μm. **F**. Schematic model depicting SKOR1 as a mediator of FER-dependent Smad2/3 signaling pathways. Error bars denote SD; * P<0.05, ** P<0.01, **** P<0.0001, ns indicates non-significant; one-way ANOVA.

## Discussion

Because SKOR1 expression has been predominantly observed in brain tissues, most studies have focused on its role during neuronal development(Arndt et al., 2007). Now, our study identifies SKOR1 as a direct FER substrate *in vitro* and an important regulator of breast cancer growth and metastasis formation *in vivo*. We propose that SKOR1 plays a key role in promoting cell proliferation and migration, thereby driving tumor progression and metastasis formation. Interestingly, SKOR1 expression has been associated with migration of other cell types. SKOR1 is expressed in Purkinje cells throughout all stages of embryonic development and in adulthood, with particularly high levels noted in the migratory precursors of Purkinje cells(Arndt et al., 2007). In early stages of wound healing, SKOR1 expression is upregulated in fibroblasts, where it may control cellular migration by affecting F-actin dynamics and/or organization(Arndt et al., 2011). Elevated SKOR1 levels are sustained in the fibro-proliferative diseases keloid scars and skin sclerosis^28^, possibly contributing to the pathogenesis of these diseases by promoting the migration of dermal fibroblasts into the wound site, reorganizing the collagen structure that is deposited by these fibroblasts and controlling collagen contraction(Arndt et al., 2011). In line with our data showing that SKOR1 loss induces stress fiber and FA formation and impairs migration, overexpression of SKOR1 in fibroblasts promotes cell motility by driving F-actin and FA complex redistribution at the cell periphery(Arndt et al., 2011). Based on these observations and our findings, SKOR1 appears to play a critical role in regulating cytoskeletal organization to control cell morphology and promote cell motility(Arndt et al., 2011).

Here, we propose that SKOR1 acts downstream of FER, because SKOR1 loss of function phenotypes are virtually identical to FER kinase inactivation(Ivanova et al., 2013; Sangrar et al., 2007). In previous studies we determined that FER regulates actin dynamics and FA distribution via endosomal recycling of growth factor receptors and cell adhesion molecules, and that dysregulation of membrane trafficking upon FER depletion greatly impairs the migratory and invasive behaviour of breast cancer cells(Ivanova et al., 2013). Because SKOR1 belongs to a well-known family of major regulators of TGFβ signaling, a pathway for which endosomal recycling of TGFβ receptors is essential(Yakymovych et al., 2017), we decided to further study the role of SKOR1 in TGFβ signaling promoting FER-dependent cytoskeletal organization and invasion in TNBC. In early endosomes, signaling-promoting factors such as SARA (Smad Anchor for Receptor Activation) support Smad2/3 and TGFβ receptor interaction, which facilitate TGFβ signaling and downstream Smad2/3 phosphorylation(Chen, 2009; Nawshad et al., 2005). Phosphorylated Smad2/3 forms a heterotrimeric complex with Smad4 that translocates to the nucleus where it associates with transcription factors and coregulators to control expression of >500 specific target genes(Tecalco-Cruz et al., 2018). Interestingly, our results show that loss of FER expression and SKOR1-Y234F substitution similarly affect (phosphorylation) levels of several growth factor receptors and downstream signaling factors, including decreased phosphorylation of TGFβ signaling mediators Smad2/3. Like SARA, Ski, SnoN, and SKOR2 are Smad-interacting proteins that regulate TGFβ signaling through simultaneous interaction with Smads(Wu et al., 2002; Tecalco-Cruz et al., 2018). Interestingly, we find that SKOR1 preferentially binds to Smad3 in TNBC cells, an observation that is in agreement with previous reports(Arndt et al., 2007; Takaesu et al., 2012). While Ski and SnoN proteins mainly localize to the nucleus to bind Smad4 and act as transcriptional corepressors through the I-loop of their SAND-domain(Nicol and Stavnezer, 1998; Nicol et al., 1999), SKOR1 shows a predominant cytoplasmic and vesicular-like localization in MM231 cells, suggesting control over Smad3 in the cytosol. Our data indicate that FER phosphorylates SKOR1 on tyrosine Y234, a residue located in the I-loop of SKOR1^25^, and that loss of function phenylalanine exchange at this site increases Smad3 binding to SKOR1. These results suggest that FER-dependent SKOR1 phosphorylation promotes dissociation of SKOR1-Smad2/3 complexes and thereby potentiates TGFβ signaling through a release of Smad2/3 molecules and subsequent recruitment by signaling-promoting factors such as SARA (Fig. 6F). Furthermore, we suggest that potentiation of TGFβ signaling through releasing Smad3 upon phosphorylation of SKOR1-Y234 by FER leads to transcriptional expression of plasminogen activator inhibitor type-1 (*PAI-1*), a known player in cancer progression(Petersen et al., 2010). *PAI-1* expression is known to induce tumor angiogenesis(Isogai et al., 2001) and promotes breast cancer cell migration through induction of F-actin-dependent formation of membrane protrusions(Chazaud et al., 2002; Liu et al., 2020). Hence, although the exact mechanism whereby SKOR regulates FA distribution and actin dynamics is still unclear, these studies suggest that SKOR1 activity can induce cytoskeletal organization and promote cell migration possibly through Smad3 signaling and *PAI-I* expression. It has also been reported that Smad3 can upregulate the expression of ubiquitin ligases that target RhoA for degradation(Yu et al., 2015). Activation of RhoA is essential for FA assembly and stress fiber formation(Aguilar-Rojas et al., 2012), features that were evident in SKOR1-depleted and SKOR1-Y234F expressing cells. These data suggest that SKOR1 loss or SKOR1-Y234F expression might cause sustained RhoA activity by decreasing Smad3 signaling, thereby inducing F-actin bundling and FA formation. Conversely, SKOR1-Y234 phosphorylation by FER could promote Smad3-dependent degradation of RhoA, leading to F-actin redistribution and disassembly of FAs to increase cell migration and invasion.

In closing, we present evidence that SKOR1 mediates FER-dependent tumor cell migration and invasion in breast cancer cells through regulation of actin cytoskeleton dynamics and FAs formation. Our data show that the SKOR1 tyrosine 234 residue, a candidate FER kinase phosphorylation site, is critical for the invasive growth of basal-type breast cancer cells. Although further studies will be needed to provide formal proof for phosphorylation of SKOR1-Y234 *in vivo*, our work substantiates FER as a cardinal tumor progression factor and advocates inhibition of FER kinase activity as a promising intervention for high-grade and basal-type breast cancers.

## Materials & Methods

### Constructs, virus generation and transduction

SKOR1 Exon 5 wild-type (WT) and Y^NULL^, in which all tyrosine residues were substituted by alanine residues, were ordered as gene blocks, subcloned in the pJET1.2/blunt cloning vector (Fermentas by Thermo Scientific, St. Leon-Rot, Germany) and inserted into the *Not*I/*EcoR*I sites of pGEX-6P-1 to introduce the GST tag. For stable knock-down (KD) of SKOR1, human SKOR1 shRNA pLKO.1-puro constructs were used (#1: 5’-CGAGCCAGATAAGGAAGACAA-3’, #2: 5’-CCTATCCAGACCAAAGGAGTA-3’). pLKO.1-TRC (shSCR) (Addgene; 10879) was used as a control. The inducible SKOR1 RNAi system was generated as described previously using the shSKOR1 #1 sequence(Schackmann et al., 2011). To generate GFP-SKOR1-expressing constructs, full length SKOR1 cDNA was obtained from transOMIC (BioCat, gene ID 390598) and was cloned into a Gateway-compatible entry vector (Thermo Fischer scientific) (pENTR-SKOR1), and subsequently recombined into destination vectors using the Gateway™ LR Clonase™ II Enzyme Mix (Invitrogen, 11791-020). Mutations were introduced in pENTR-SKOR1 following the QuikChange II XL Site-Directed Mutagenesis protocol (QuikChange; Agilent Technologies, Wilmington DE). Three silent mutations were introduced to create resistance to sh901 (SKOR1-shRes) using primers 5’-GAAACGAGGAAATCCTA**<u>C</u>**CCAGACCAAAG**<u>A</u>**AG**<u>C</u>**ATCTCCCAGCC-3’ (forward) and 5’-GGCTGGGAGAT**<u>G</u>**CT**<u>T</u>**CTTTGGTCTGG**<u>G</u>**TAGGATTTCCTCGTTTC-3’ (reverse). The Y234F mutation was introduced using primers 5’-CCGACGCCAAGTTCACGCAGCCCGA-3’ (forward) and 5’-TCGGGCTGCGTGAACTTGGCGTCGG-3’ (reverse), generating pENTR-SKOR1-shRes-Y234F. SKOR1-shRes-WT and -Y234F were then recombined into pLenti CMV-GFP (658-5) (Addgene; 17448). For transduction of the above mentioned constructs, pLV-PGK-GFP (Addgene; 19070) was included as a control. Lentivirus was produced in HEK239T cells, followed by cell transduction as described previously(Schackmann et al., 2011) and one week of puromycin selection (2 µg/ml) for stable SKOR1 KD MM231 cells. For CRISPR-based gene editing, oligo’s encoding single guide RNA sequences targeting FER (sense: 5’-AAATCCTTGGAGACTTTACG-3’ and anti-sense: 5’-CGTAAAGTCTCCAAGGATTT-3’) were annealed by heating to 95°C, followed by a gradual cool-down to room temperature. Annealed oligos were ligated into *Bbs*I-digested pACEBac1-Cas9-GFP. Left and right homology arms (LHA and RHA, respectively) were designed according to the In-Fusion cloning strategy and ordered as gene blocks (Supplementary info). Each homology arm was subcloned into the pJET1.2/blunt cloning vector (Fermentas by Thermo Scientific, St. Leon-Rot, Germany) and inserted into pUNKI-puro (kindly provided by S. Lens, University Medical Center Utrecht). LHA was first ligated into the *Cla*I/*Asc*I sites of pUNKI, followed by ligation of RHA into the *Sac*I/*Sal*I sites to generate pUNKI-puro-LHA-RHA. The puro-LHA-RHA cassette was then inserted into the gRNA-containing pACEBac1-Cas9-GFP by *Not*I restriction cloning, generating pACEBac1_FER-AS. All constructs were verified by Sanger DNA sequencing.

### Cell culture and transfection

MDA-MB-231 (MM231) cells were obtained from Cell Lines Service (Eppelheim, Germany), STR verified, and grown in DMEM growth medium (Invitrogen, 11039-047), supplemented with 1% penicillin-streptomycin (Invitrogen, 15070-063) and 10% fetal bovine serum (FBS) (Invitrogen 16050-122). Cells were cultured at 37°C with 5% CO2. MM231 FERiKD cells have been generated previously(Ivanova et al., 2013). MM231 cells were transfected with pACEBac1_FER-AS using FuGene HD according to manufacturer’s instructions. Two days after transfection, cells were treated with 2 µg/ml puromycin for one week, followed by single-cell expansion. Positive FER^ASKI^ clones were selected by PCR and sequencing with primers 5’-TGAGGGAAGGCTTTACTCGTT-3’ (forward) and 5’-TCCTTGGAGACTTTACGAGGAG-3’ (reverse). MM231 FER^ASKI^ cells were treated with 1 µM NM-PP1 (Calbiochem) for 3 days. Dox-inducible cell lines were treated for five days with 2 µg/mL Doxycycline (Sigma-Aldrich, D9891-1G), refreshed on day 3.

For 3D assays, MM231 cells were added to Cultrex Basement Membrane Extract (BME) (Trevigen; 3533-005-02) at a density of 500 cells/100 µL BME. Droplets of 25 µL were added to flat bottom, optical plastic 24-well plates (Corning, Tewksbury, USA). Plates were incubated for 45 min at 37°C to allow the BME to solidify, after which 500 µL normal growth medium was added. Cells were cultured for 7 days at 37°C.

### 3D Morphology assessment

Brightfield images were acquired by using a 10× objective on an EVOS M5000 Imaging System (Thermofisher). At least 5 images were acquired per chamber well, and at least two wells were imaged per condition. Each image was segmented by individually optimizing the OrganoSeg(Borten et al., 2018) parameters manually until a suitable segmentation was achieved. Invasiveness was inferred using ‘Solidity’ parameter (*Invasiveness=1-Solidity*) reported in each spheroid caption.

### Expression of recombinant proteins and in vitro kinase assays

Protein expression in the *E. coli* strain Rosetta was induced by incubation with 0.2mM Isopropyl β-D-1-thiogalactopyranoside (IPTG) for 16h at 18°C. Recombinant proteins were extracted from bacterial pellets by adding lysis buffer (10mM EGTA, 10mM EDTA, 0,1% Tween 20, 250mM NaCl, 5mM DTT) supplemented with 0.325 mg/mL lysozyme and a cocktail of protease inhibitors (Roche, Complete™ EDTA free). Samples were sonicated, centrifuged and coupled to glutathione-Sepharose 4B beads (Amersham Biosciences), followed by elution with elution buffer (100mM Tris (pH8.0), 30mM GSH and 75mM KCl). Eluates were treated with sample buffer (375 mM Tris-HCl pH 6.8; 25% glycerol; 12.5% β-mercapto ethanol 10% SDS: 0.025% bromophenol blue) and boiled. Proteins were separated on 15% SDS-PAGE gel and stained with Instant Blue. Recombinant analogue-sensitive (AS) or kinase-dead (D742R) GST-FER were incubated with WT or Ynull GST-SKOR1, or GST-Cortactin as substrate. Each reaction was performed in 25 µL kinase buffer (10 mM MnCl2, 20 mM Tris-HCl pH 7.5, 0.1 mM sodium orthovanadate) supplemented with 250 µM N-benzyl (Bn)-ATPγS (B 072-05, BioLog), with or without 10 µM NM-PP1. After 30 min incubation at 30°C, reactions were terminated by addition of 2.5 mM EDTA were subsequently incubated with 2.5 mM p-Nitrobenzyl mesylate (Epitomics, Burlingame, CA) for 2h at room temperature. Reactions were stopped by adding sample buffer.

### Immunoblotting

Proteins were extracted from cells by scraping in sample buffer and lysed for 10 min on ice, followed by 10 min of boiling. Protein extracts and *in vitro* kinase reaction products were separated by SDS-PAGE electrophoresis and blotted onto PDVF membrane. Following 1 h blocking with 5% bovine serum albumin (BSA) in tris-buffered saline (TBS) 0.1% Tween20, the membrane was incubated with primary antibodies in blocking buffer overnight at 4°C. After washing three times with PBS-Tween 0.1%, membranes were probed with either horseradish peroxidase-conjugated secondary antibodies (DAKO; 1:10,000) or IRDye 680- and 800-conjugated secondary antibodies (LI-COR; 1:5,000) for 1 h at room temperature and visualized using Enhanced Chemo-Luminescence (ECL) (GE Healthcare) or Typhoon Biomolecular Imager (GE Healthcare), respectively. The following primary antibodies and dilutions were used for western blotting: mouse anti-GAPDH (Millipore, Mab374; 1:1,000), rabbit anti-Thiophosphate ester (Abcam, ab92570; 1:5,000), mouse anti-GST (Santa Cruz, sc-138; 1:1,000), mouse anti-GFP (Santa Cruz, sc-8334, 1:500), rat anti-GFP (3HG) (Chromotek, 029762; 1:1,000), goat anti-Akt (Santa Cruz, sc-1618; 1:1,000)mouse anti-pY20 (BD Biosciences, 610011; 1:1,000), rabbit anti-LBXCOR1 (Sigma, SAB2105374; 1:500), mouse anti-Smad2/3 (C-8) (Santa Cruz, sc-133098; 1:1,000), rabbit anti-phospho-EGFR Tyr1173 (Cell Signaling Technology, 4407, 1:500).

### 3D invasion assay, BrdU incorporation and immunofluorescence

Invasion and proliferation were assessed by incubating cells in BME (3D) with 10 µm BrdU for 2 h. Cells were fixed with 4% paraformaldehyde (PFA) 1% Glutaraldehyde (Sigma, G5882), followed by 1% NaBH4 treatment for 30 min. Then cells were washed with PBS before 2M HCl treatment for 80 min, Fixed cultures (3D) were blocked for overnight in 5% goat serum 0.3% Triton in PBS and incubated with 10 µL/mL Alexa 647-conjugated anti-BrdU antibody (BD biosciences, 560209) or anti-cleaved Caspase 3 (Cell Signaling, 9661s, 1:250) in 1% BSA 0.3% Triton in PBS overnight. Structures were washed three times and then probed with secondary antibodies (when applicable), Alexa 568-conjugated Phalloidin (ThermoFisher; A12380; 1:200) and DAPI in 1% BSA 0.3% Triton in PBS overnight.

For 2D assays, cells were grown on 12 mm coverslips and fixed for 30 min using 4% PFA. Fixed cells were permeabilized with 0.1% Triton X-100 in PBS for 3 min, blocked with 5% BSA for 30 min and incubated with primary antibodies in 1% BSA in PBS overnight at 4°C. Cells were washed three times and then probed with secondary antibodies and Alexa 568-conjugated Phalloidin (ThermoFisher; A12380; 1:200) in 1% BSA for 2 h. After three washes in PBS, cells were stained with 2 µg/ml DAPI (Sigma; D9542) in 1% BSA for 5 min. Again, cells were washed thrice with PBS before mounting using ProLong Diamond Antifade (Invitrogen). Primary antibody used was rabbit anti-phospho-Paxillin (Life Technologies, 447226; 1:200). Secondary antibody used was goat anti-rabbit Alexa-647 (Molecular Probes by Invitrogen).

An inverted Carl Zeiss LSM 700 Laser Scanning Microscope with a Plan-Apochromat 63x/1.40 Oil DIC M27 or with a LD Plan-Neofluar 40x/0.6 Korr M27 objective was used for imaging. Confocal images were analyzed using ImageJ Software.

### Co-Immunoprecipitation

GFP-tagged proteins were immunoprecipitated using a GFP-Trap system (Chromotek, gta-10) as described(Ven et al., 2016). After washing with lysis buffer, bound and non-bound proteins were eluted in sample buffer, boiled, and analyzed by western blot.

### Cell migration assay

MM231 SKOR1iKD cells were induced for 5 days with or without Dox, and MM231 FER^ASKI^ cells were treated for 3 days with NM-PP1 or DMSO (control), before plating them on 24-well plastic-bottom plates (Corning, Tewksbury, USA). Cells were incubated with 200 nM SiR-DNA (Spirochrome) for 7-8 hours before imaging and were imaged every 10 min for 16 h using a Carl Zeiss Cell Observer widefield microscope with an EC Plan-Neofluar 5x/0.16 M27 objective. During imaging, cells were kept in complete DMEM medium (with or without DOX, or with DMSO or NM-PP1) under normal growth conditions (37°C, 5% CO^2^). Cell migration was quantified using the Imaris for Tracking software (Bitplane, Oxford Instruments, UK).

### Quantitative real-time PCR

Total RNA from the samples was extracted using RNeasy Plus Mini Kit (Qiagen, 74104) following the manufacturer’s guidelines. cDNA synthesis was performed according to iScript cDNA Synthesis Kit (Bio-Rad). qPCR reactions were performed using FastStart Universal SYBR Green Master mix (Roche, 4913957001) and Bio-Rad CFX96 touch Real-Time PCR detection system (Bio-Rad). mRNA levels were normalized to their corresponding GAPDH expression levels. Knock-down efficiency was quantified using the Pfaffl method(Pfaffl, 2001). Used oligonucleotide primer pairs: SKOR1 forward 5’-CCACGAGCCAGATAAGGAAG-3’, SKOR1 reverse 5’-CCATTTGTTCCAGGAGCAGT-3’, PAI-I Forward 5’-TCTTTGGTGAAGGGTCTGCT-3’, PAI-I Reverse 5’-CTGGGTTTCTCCTCCTGTTG-3’, GAPDH forward 5’-TGCACCACCAACTGCTTAGC-3’ and GAPDH reverse 5’-GGCATGGACTGTGGTCATGAG-3’.

### Colony formation assays

MM231 SKOR1iKD cells were cultured with or without Dox for 5 days, and FER^ASKI^ cells were treated with DMSO or NM-PP1 for 3 days before seeding them at a density of 7,500 cells into each well of a 24-well plate and culturing them for 3 days. Cells were fixed using 10% Glutaraldehyde in normal growth medium for 10 min, stained using 0.1% crystal violet (Sigma) for 30 min while agitating, washed with water and dried overnight. Acetic acid 10% (v/v) was used to elute incorporated crystal violet for 30 min while agitating, after which the solution was quantified with a spectrophotometer at 590 nm (Bio-Rad).

### Reverse protein phase array (RPPA)

MM231 SKOR1iKD cells were cultured with or without Dox for 5 days, and FER^ASKI^ cells were treated with DMSO or NM-PP1 for 3 days. Cell extracts were prepared in RIPA (50 mM Tris-HCl pH 7.4, 150 mM NaCl, 1 mM EDTA pH 8.0, 1 mM EGTA, 1% Triton X-100, 0.1% SDS, 10 mM Glycerolphosphate, protease inhibitor cocktail (Complete, Sigma) and phosphatase inhibitor (phosSTOP, Roche). RPPA was performed as described(Teo et al., 2018).

### Mouse studies

Recipient female RAG2-/-;IL-2Rγc-/- immunodeficient mice (Envigo) were orthotopic transplanted with luciferase-expressing MM231 cells (FER^ASKI^ or SKOR1iKD), using a 50-μl Hamilton syringe (Hamilton, Bonadur, Switzerland). Mice were anesthetized using isoflurane (IsoFlo; Le Vet Pharma). Burprenorphine (0.1mg/kg) was injected subcutaneously as analgesic treatment. After a recovery period of 2 weeks, mice were anesthetized with IsoFlo, injected i.p. with 225 μg/g body weight n-luciferin (potassium salt; Biosynth AG) and imaged on a Biospace Φ bioluminescence imager (Biospace Lab). Tumor growth was measured using a digital pressure-sensitive caliper (Mitutoyo) on a weekly basis. Treatment started when tumors reached a volume of 50-100mm^3^. To study *in vivo* inactivation of FER, FER^ASKI^ mice were switched to water containing DMSO (solvent) or NM-PP1 (25µM, Merck) *ad libitum*. Drinking water was refreshed twice a week. To deplete SKOR1 in cancer cells, SKOR1iKD mice were switched from standard diet to doxycycline containing chow (200mg/kg; A155D70201; Ssniff, Bio services) *ad libitum*. Mice were euthanized if mammary tumor reached a size of 1000mm^3^, in case of severe discomfort or when bioluminescence imaging revealed metastases.

### Studies approval

All animal experiments were performed in accordance with local, national and European guidelines under permit AVD115002015263 issued by The Netherlands Food and Consumer Product Safety Authority (NVWA) of the ministry of Agriculture, Nature and Food.

### Statistical analysis

Statistical analyses were performed using IBM SPSS Statistics (SPSS Inc., Chicago, IL, USA). The Kaplan-Meijer method was used for cumulative survival analysis; a two-way mixed model analysis of variance was used to evaluate differences in tumor volume. For RPPA data, Z-scores were clustered (Cluster software version 3.0) and visualized in heatmaps (Java Treeview). All other data were analyzed using one-way ANOVA or unpaired t-test methods in Prism (Graphpad software version 8.0).

## Supporting information

Table S1 - MM231 FERiKD RPPA list of significant substrates

Table S2 - SUM149PT FERiKD RPPA list of significant substrates

Table S3 - MM231 SKOR1recon RPPA list of significant substrates

## Online supplemental material

Fig. S1 shows how FER^ASKI^ cells were generated and validated. Fig. S2 shows that FER regulates key signaling pathways in MM231 breast cancer cells. Fig. S3 shows that FER regulates key signaling pathways in SUM149PT breast cancer cells. Fig. S4 shows that SKOR1 is necessary for proliferation of MM231 breast cancer cells. Table S1 lists RPPA significant substrates upon FER depletion in MM231 cells. Table S2 lists RPPA significant substrates upon FER depletion in SUM149PT cells. Table S3 lists RPPA significant substrates upon SKOR1^Y234F^ reconstitution in MM231 cells.

## Acknowledgements

We thank Peter ten Dijke for reagents and comments on the manuscript. This research was supported by KWF Kankerbestrijding (UU2014-7201), The Netherlands Organization for Scientific Research (NWO-TOP 02007), and the European Union’s Horizon 2020 FET Proactive program under the grant agreement No. 731957 (MECHANO-CONTROL).

## Author contributions

Conceptualization, S.T. and P.W.B.D.; methodology, S.T. and P.W.B.D.; investigation, E.B., S.T., L.S., E. B., C.O., and L. E.; formal analysis, S.T., L.S., M.R., E. B. and L.E.; resources, M. H. and P. t. D.; writing—original draft, L. S. and S.T.; writing—review and editing, V.G. B, S.T. and P.W.B.D; funding acquisition, P.W.B.D; supervision, S. T. and P.W.B.D.

