## Supplementary figures and images for "SKOR1 mediates FER kinase-dependent invasive growth of breast cancer cells"

### Supplementary Figure 1

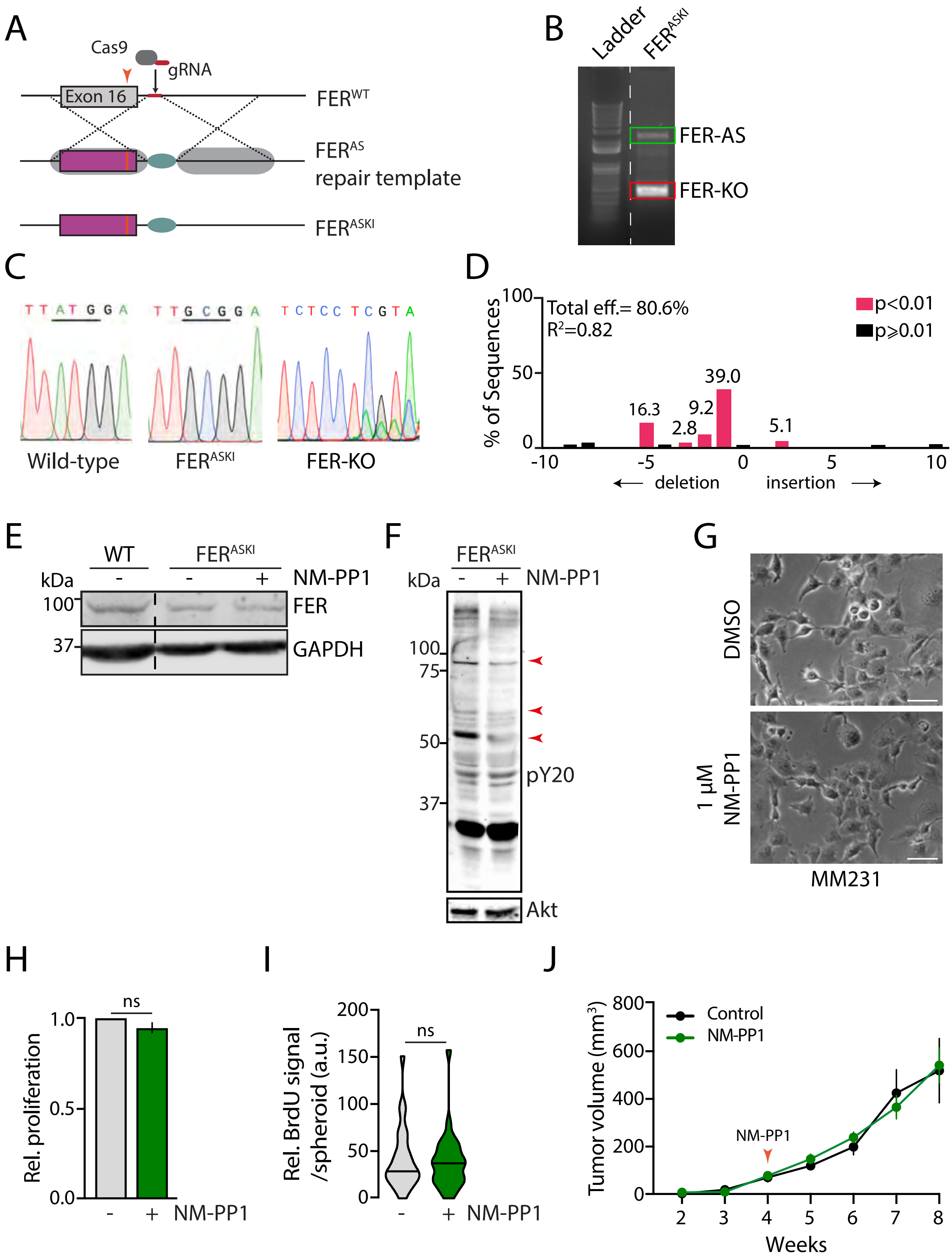

### Supplementary Figure 2

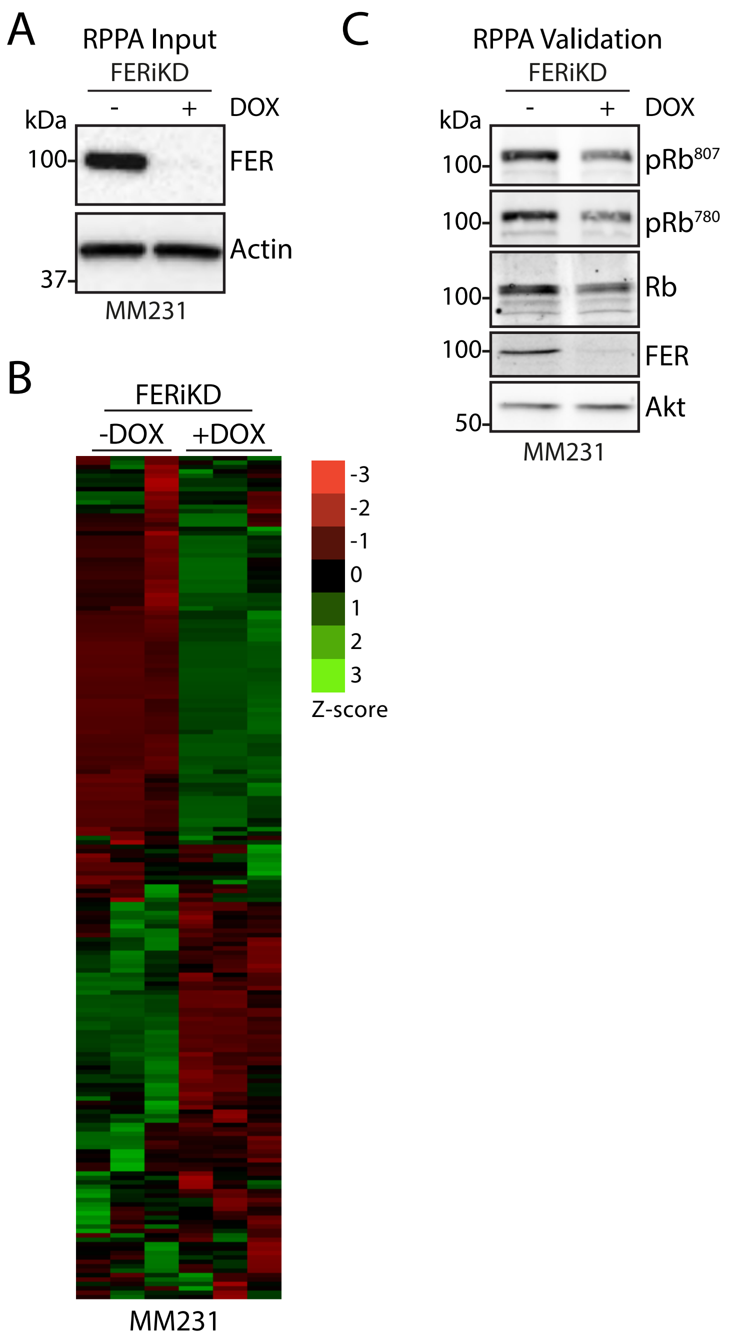

### Supplementary Figure 3

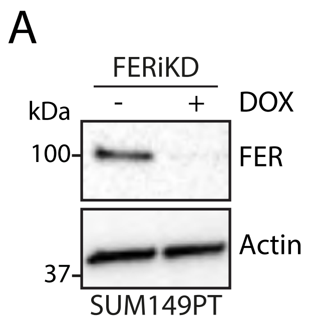

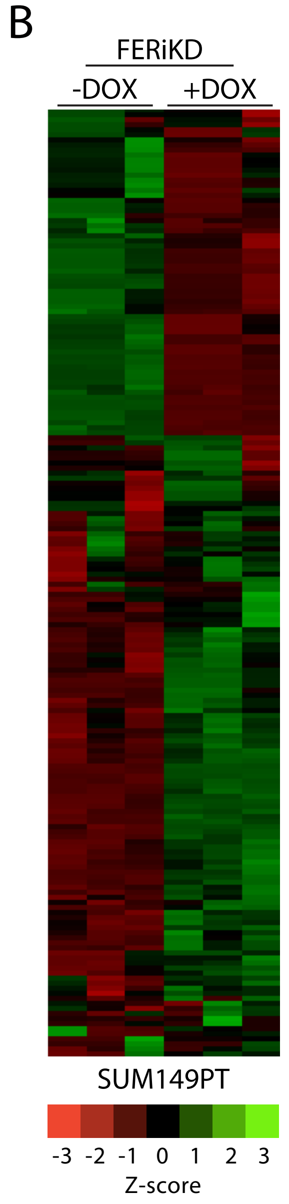

### Supplementary Figure 4

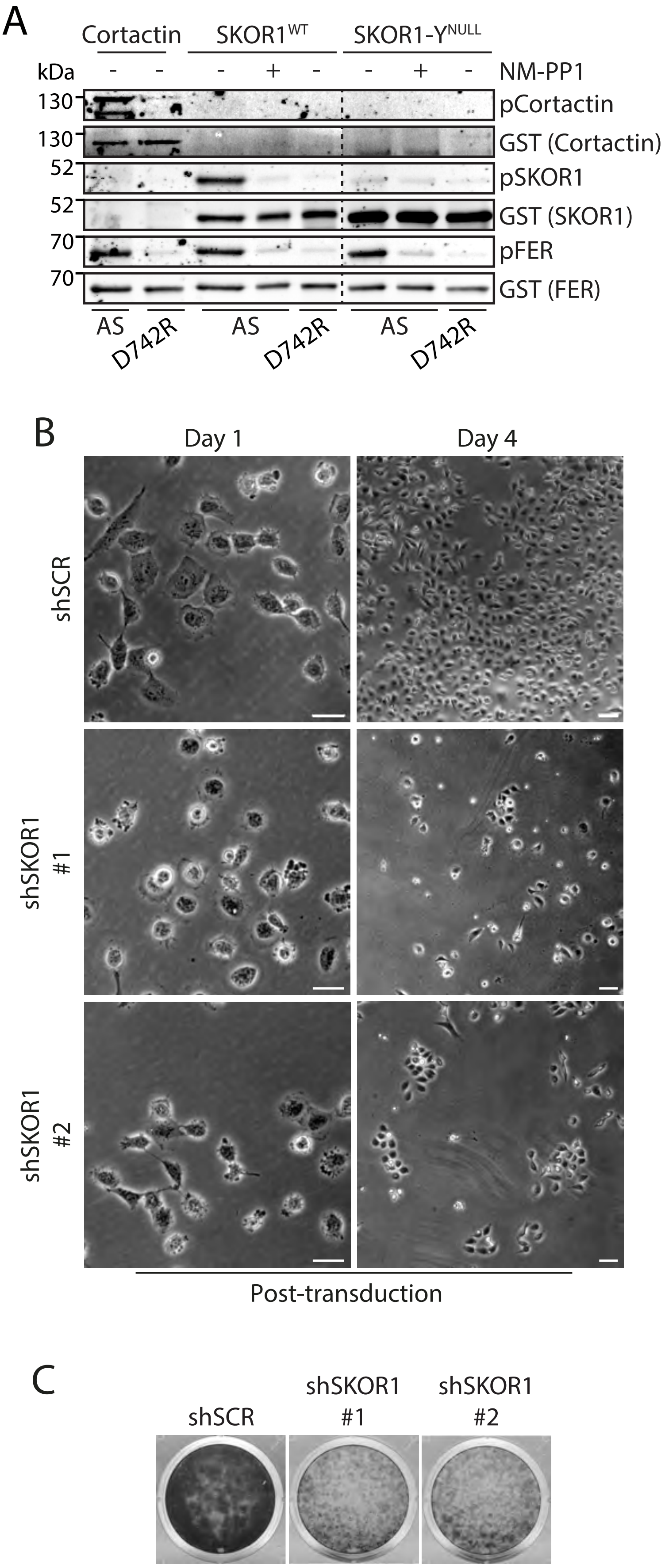
